# The equivalent arc ratio for auditory space

**DOI:** 10.1101/134700

**Authors:** W. Owen Brimijoin

## Abstract

The minimum audible movement angle increases as a function of source azimuth. If listeners do not perceptually compensate for this change in acuity, then sounds rotating around the head should appear to move faster at the front than at the side. We examined whether judgments of relative amounts of acoustic motion depend on signal center angle and found that the azimuth of two signals strongly affects their point of subjective similarity for motion. Signal motion centered at 90° had to be roughly twice as large as motion centered at 0° to be judged as equivalent. This distortion of acoustic space around the listener suggests that the perceived velocity of moving sound sources changes as a function of azimuth around the head. The “equivalent arc ratio,” a mathematical framework based on these results, is used to successfully provide quantitative explanations for previously documented discrepancies in spatial localization, motion perception, and head-to-world coordinate transformations.

## 1 Introduction

The binaural cues that are used to construct our internal representation auditory space are interaural time difference (ITD) and interaural level difference (ILD). Both of these arise from the physical structure of the head, as sounds arrive at the ear closest to the sound source both earlier and at a higher level than at the further ear. Measurements of these binaural cues as a function of sound source angle demonstrate that they change most rapidly at the front of a listener [1, 2]. If the internal representation of space were based purely on these cues, then listeners would have increased spatial resolution near the sagittal plane. Indeed this is well supported: both the threshold measurements of minimum audible angle (MAA) and minimum audible movement angle (MAMA) are known to change as a function of source azimuth [3, 4].

The study described here examined a potential consequence of this representation of auditory space: namely that if listeners do not perceptually compensate for it, then the expansion of resolution at the front and contraction at the side would dictate that a sound rotating at a constant angular velocity around the head would not appear to do so, but would instead appear to move faster at the front than at the side. Correspondingly, listeners turning their heads at a constant velocity would experience an angle-dependent change in apparent source movement. The literature on auditory motion is unclear on this subject and to our knowledge no studies have directly examined the perceived difference in auditory motion at different angles and directions relative to the head. That said, a number of curious discrepancies in spatial auditory perception have been described over the years. Some of these have been classed as ‘incomplete coordinate transformations’ [5], suggesting that a person’s head angle may affect the direction from which they perceive a sound to emanate. Similarly, studies have reported discrepancies in listeners’ subtraction of their own active head movements from the movement of the auditory scene [6, 7].

Although the latter two studies used Bayesian inference as a description of their observations, no mathematical framework has been suggested to account for these findings. We propose instead that these and a number of other discrepancies in spatial hearing could be explained via an angle-dependent distortion in apparent sound location and motion. The underlying hypotheses to be tested can be more formally stated as follows: first, that relative velocity judgements should change as a function of azimuth, and second, that a quantitative model based on any observed expansion of space should capture static phenomena such sound source eccentricity overestimation [8-11] and dynamic phenomena like inconsistent self-motion subtraction [6, 7]. We further argue that the MAMA is not simply a measure of acuity, rather it underlies a constant perceptual unit, changing in absolute magnitude across space. One could argue that as a threshold measurement, roughly 1-2 degrees at the front of a listener may be considered equivalent to roughly 4 degrees at the side of the listener in that these are the minimal amounts of motion required for a listener to change their judgement from no motion to ‘some motion.’ Whether a similar azimuth-dependent expansion in the perception of auditory space exists at suprathreshold levels has never been directly demonstrated.

We term the proposed relationship of equal perceptual units across angle the “equivalent arc ratio” for auditory space, borrowing terminology from the equivalent rectangular bandwidth [12]. Here we quantify perceptual expansion by measuring the dependence of judgements of relative sound-source motion on the angles with respect to the head from which the signals arrive. The consequences of such a nonlinear representation of space are discussed, as is the potential for ‘hyper-stable’ virtual acoustics. Finally, successful quantitative comparisons are made between the predictions of a proposed mathematical framework and the results of a number of previously published studies.

## 2 Results

### 2.1 Relative motion judgements

Listeners were asked to make a judgement about which of two signals, a reference and a test signal, “moved more.” The reference always moved 20° and the test moved less, the same, or more. Both the test and reference signals could be centered at 0°, 45°, or 90°, and movement direction and the order of presentation and condition were fully randomized. Across all conditions we found that the center azimuth of both test and reference signals strongly affected subjects’ comparison of relative extents of motion, as expressed by a change in response as a function of test excursion. We found main effects of test excursion, test azimuth, and reference azimuth (F(1,6) = 338.11, p < 0.05, F(1,2) = 71.34, p < 0.05, and F(1,2) = 97.74, p < 0.05, respectively), an interaction between reference azimuth and test excursion (F(2,12) = 3.99, p < 0.05), an interaction between test azimuth and test excursion (F(2,12) = 34.24, p < 0.05), and a 3-way interaction between reference azimuth, test azimuth, and test excursion (F(2,24) = 1.92, p < 0.05). The only insignificant comparison in the test was found in the interaction between test and reference azimuths (F(2,4) = 0.76, p = 0.55).

Figure 1a illustrates this phenomenon for reference signals centered at 0°. Test signals centered at 0° (orange line) were judged to move the same amount as the 20° reference motion when the test signals also moved by 20°. Thus the point of subjective equality (PSE) for this condition was roughly 20/20 = 1. The smooth psychometric function confirms that listeners were capable of making judgements of relative motion, which is also reflected in the significant main effect of test excursion. On the other hand, test signals centered at 45° (green line) had to move by about 25° to be judged as moving the same amount as the 20° movement of the reference signal at the front. The range of movement angles we used was not sufficient to estimate how much a 90° test signal (blue line) had to be moved to reach the PSE with a reference at 0°, but the required excursion was likely much greater than 35°. In Figure 1b it can be seen that when compared to reference signals at 45°, test signals at 0° had to be moved significantly less to be perceived as moving the same amount, and test signals at 90° had to be moved significantly more. Finally, reference signals at 90° (Figure 1c) showed a pattern of expansion that was roughly inverted as compared to Figure 1a. Here it can be seen that motion at 0° had to be roughly half as large to be judged as equivalent to motion at 90°.

**Figure 1.**
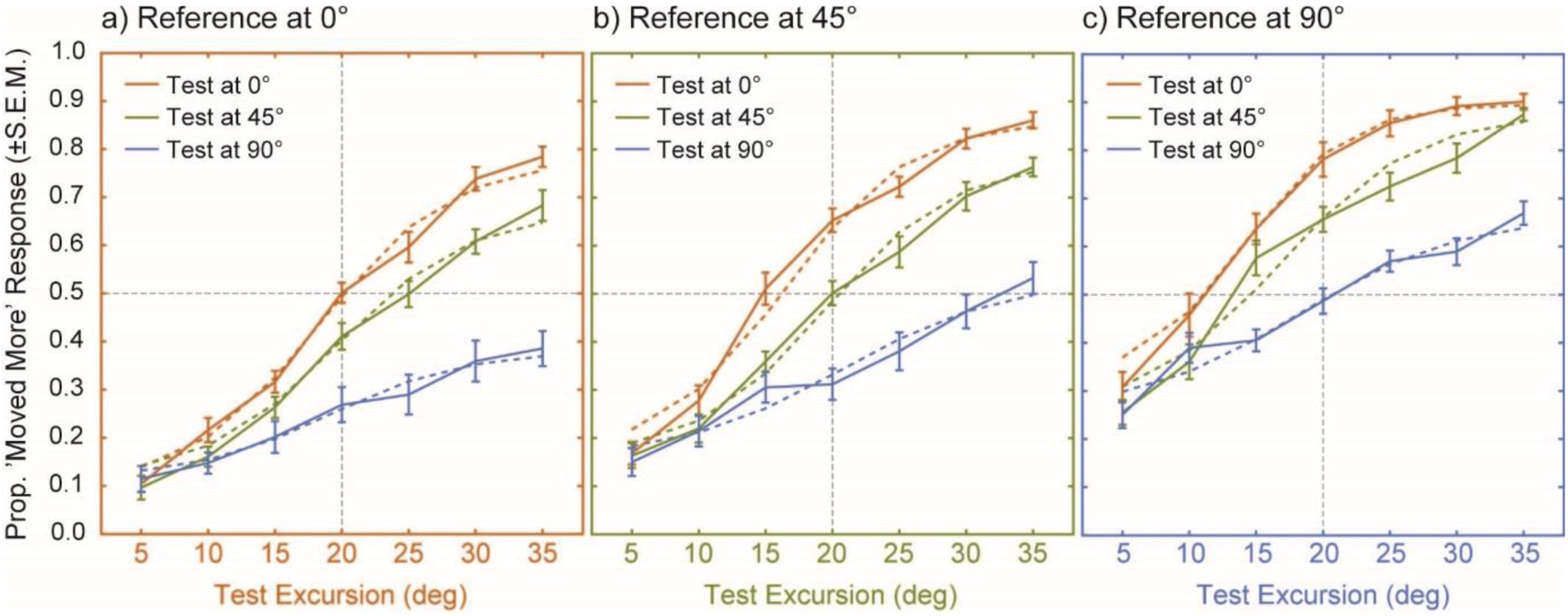
Psychometric functions for motion comparisons and points of subjective similarity (PSE) for motion for moving signals centered at 0°, 45°, and 90°. A) For conditions with a reference signal at 0°, the psychometric function for test signals also at 0° crossed the PSE at 20 degrees (orange line), whereas test signals at 45° and 90° (green and blue, respectively) had to move more to be judged as moving the same amount (rightward shift in the curves). B) Compared to 45° reference signals, 0° test signals had to be moved less (orange) and 90° signals more (blue) to be judged as moving the same amount. C). References of 90° required less motion to be judged the same as both the 0° (orange) or the 45° (green) test signals.

### 2.2 Points of subjective equality for motion

PSE ratios were drawn from the individual logistic fits to the data (the *mean* of said fits are plotted with dotted lines in Figure 1). The point at which the logistic fit crossed 0.5 probability was taken for each condition for each listener and divided by the reference motion PSE. PSE ratios were also computed for inverted pairs (i.e., the test/reference ratio 0/90 is accompanied by 1/the test/reference ratio 90/0). A scatter plot of these PSE ratios, plotted as a function of the absolute difference between the test and reference angles, is shown in Figure 2. The use of ‘comparison’ in the legend is due to the mix of normal and inverted pairs (where reference and test are used interchangeably). When the test and reference motions were identical the PSE ratios were clustered around 1, albeit with a large degree of intersubject variability. PSE ratios for a difference of 45° were on average larger than 1.0, and ratios for a 90° difference were still larger, reaching a value of roughly 2. It should be noted that the triangle symbols in the plot represent measurements in which the test signals at maximum excursion were still not judged to be moving by the same amount as the 20° reference motion. The true values for these data points cannot be reliably estimated as the psychometric functions in question did not cross the PSE, but examining the individual data and logistic fits makes it clear that the values are likely to be substantially larger than 2. The Pearson correlation coefficient between the PSE ratio and the difference in test/reference angle is R^2^ of 0.43.

**Figure 2.**
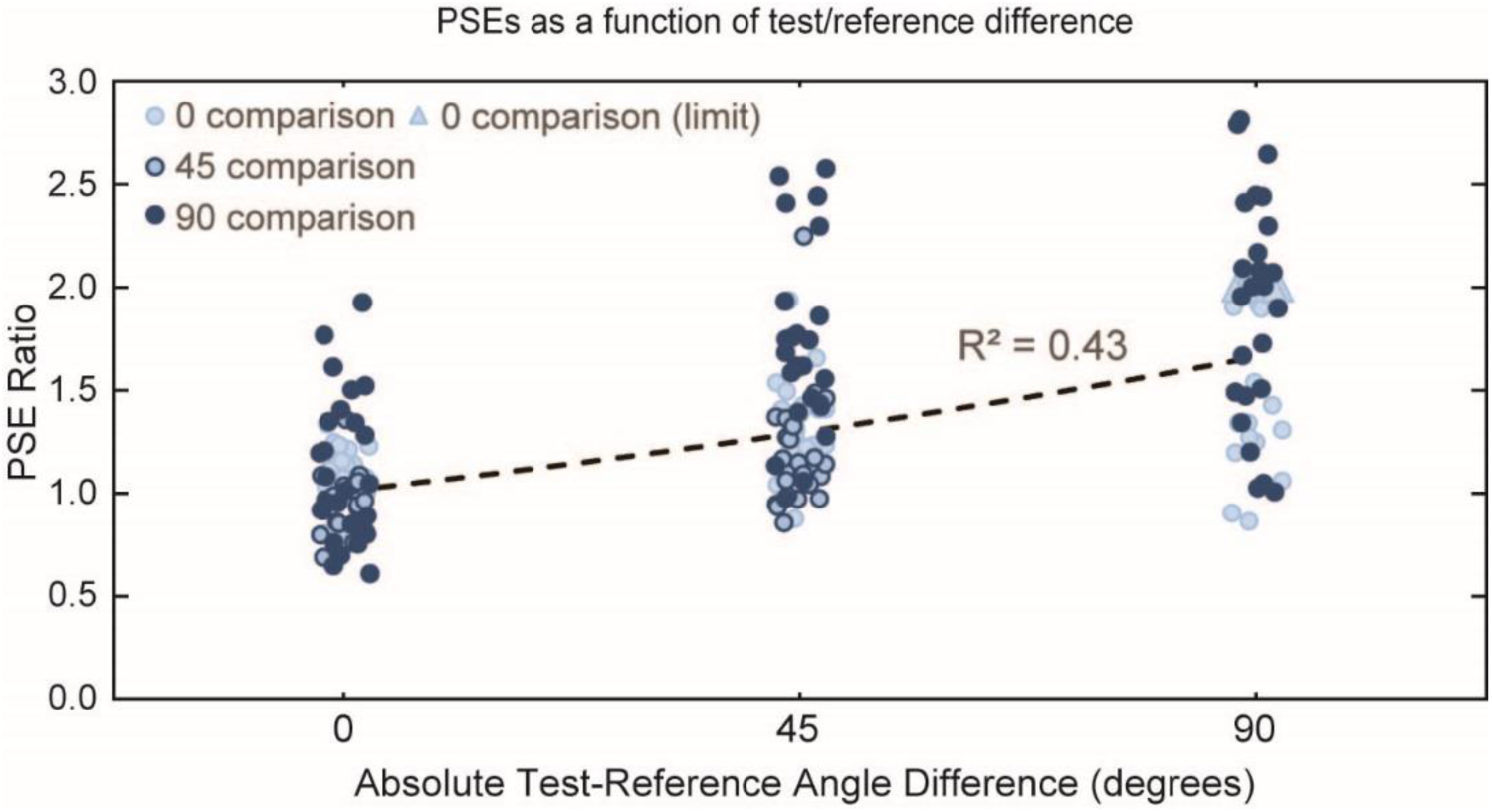
Scatter plot of PSE ratios showing an expansion of auditory space. All x values are jittered for visibility. PSE ratio increases as a function of the absolute difference between test and reference angles. The different symbols represent actual angle comparisons, some values for which were inverted from test/reference to reference/test. Triangle symbols represent measurements in which the test signals at maximum excursion were still not judged to be moving by the same amount as the 20° reference motion.

## 3 Discussion

### 3.1 The non-uniformity of acoustic motion

The observed changes in the apparent amount of motion across azimuth are not subtle, making it somewhat surprising that this effect has not been previously reported. Across all conditions we found that the relative azimuth of two signals strongly affects their point of subjective similarity for motion. Roughly speaking, 20° of motion at the front of the listeners is treated equivalently to 40° of motion at the sides. This difference in PSE ratio over azimuth, which we will refer to as the equivalent arc ratio from here onward, represents a perceptual expansion of space at the front and a contraction at the sides. On one level, the equivalent arc ratio could be interpreted as a simple relationship between acuity and perception, but this belies two perceptual consequences. One consequence is that a sound rotating at a constant angular velocity around the head would appear to accelerate towards the front of the listener, and decelerate towards the side. The second consequence is that – from the perspective of a moving listener –the acoustic world not appear stable as the head turns. Instead signals at the front should appear to counter-rotate as the listener turns, and signals at the side should seem to be slightly dragged along with the listener’s rotation.

### 3.2 Distortion in acoustic location

There are two possibilities for reconciling the observed change in perceived motion as a function of angle with our current understanding of sound localization. The first requires a disassociation between movement and location; in this case the apparent location of a signal at the end point of a movement would have to be different from its apparent position(s) during the movement. There is evidence in the visual system of just such disassociation [13]. It is conceivable that a similar process occurs in the auditory system, but the disassociation in the visual system is thought to arise from specialized motion-sensitive neurons in the middle temporal visual and medial superior temporal areas [14], brain regions known for motion selectivity [15, 16]. Motion-specific processing has been observed in posterior auditory cortex [17], however, there remains little physiological evidence for auditory neurons that exhibit motion selectivity while being agnostic to spatial location.

If we assume, on the other hand, that auditory motion and spatial location are intrinsically linked with each other, then the second possibility is that both the motion and the perceptual location of *static* sound sources would be subtly distorted as a function of head angle. This framework prevents any jump in perceived location after a movement (as would be found above), but requires that listeners mislocalize sound sources. The function and its constants were chosen so that its slope at 0° and its slope at 90° were related to each other in the same manner as the equivalent arc ratio between these two angles. We used a hyperbolic tangent (Equation 1) because it is readily invertible, although one could in principle also use a sine-expansion, or some other mathematical construct.

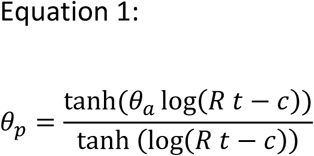

where all angles are degrees / 90 (including *Θ*_a_), ln is the natural logarithm, t is a constant equal to 7.08, c is a constant equal to 5.97, R is the ratio between the PSE at 90° and at 0°, *Θ*_a_ is the actual position of a signal, and *Θ*_p_ is the perceived position of that signal. The constants t and c were empirically derived (using Matlab’s fminsearch function) to ensure that the ratio between its 20° slope (the amount of reference motion) at 0° and at 90° was closest to the ratio R between the PSE at 0° and at 90° over a reasonable range of values of R.

For *Θ*_a_ angles larger than 0 and less than 90, the values of *Θ*_p_ generated by Equation 1 imply that static acoustic targets would be perceived at larger eccentricities than they truly are. Precisely such a phenomenon has been repeatedly demonstrated in the literature, as listeners have been shown to regularly overestimate the angle of sound sources [11], particularly when fixating at the front and using a laser pointer to indicate direction. Equation 1 provides a reasonable fit to the overestimation of source angle measured in at least three separate laser-pointer studies [8-10] (laser pointing being the most comparable task condition it does not involve a head movement). The data from the most relevant portions of these studies are plotted in Figure 3 alongside predictions from the model. The predictions are plotted as the difference between perceived and actual locations (*Θ*_p_ - *Θ*_a_), and these values fall well within the range of the data from the three studies. Physiological data on this subject are somewhat limited, but predictions of a neural network trained on spike data from cat primary auditory cortex also show a characteristic overestimation of target position that roughly follows the predicted pattern [18]. The magnitude of this overestimation, however, is far larger than has been observed behaviorally or predicted by the current mathematical framework.

**Figure 3.**
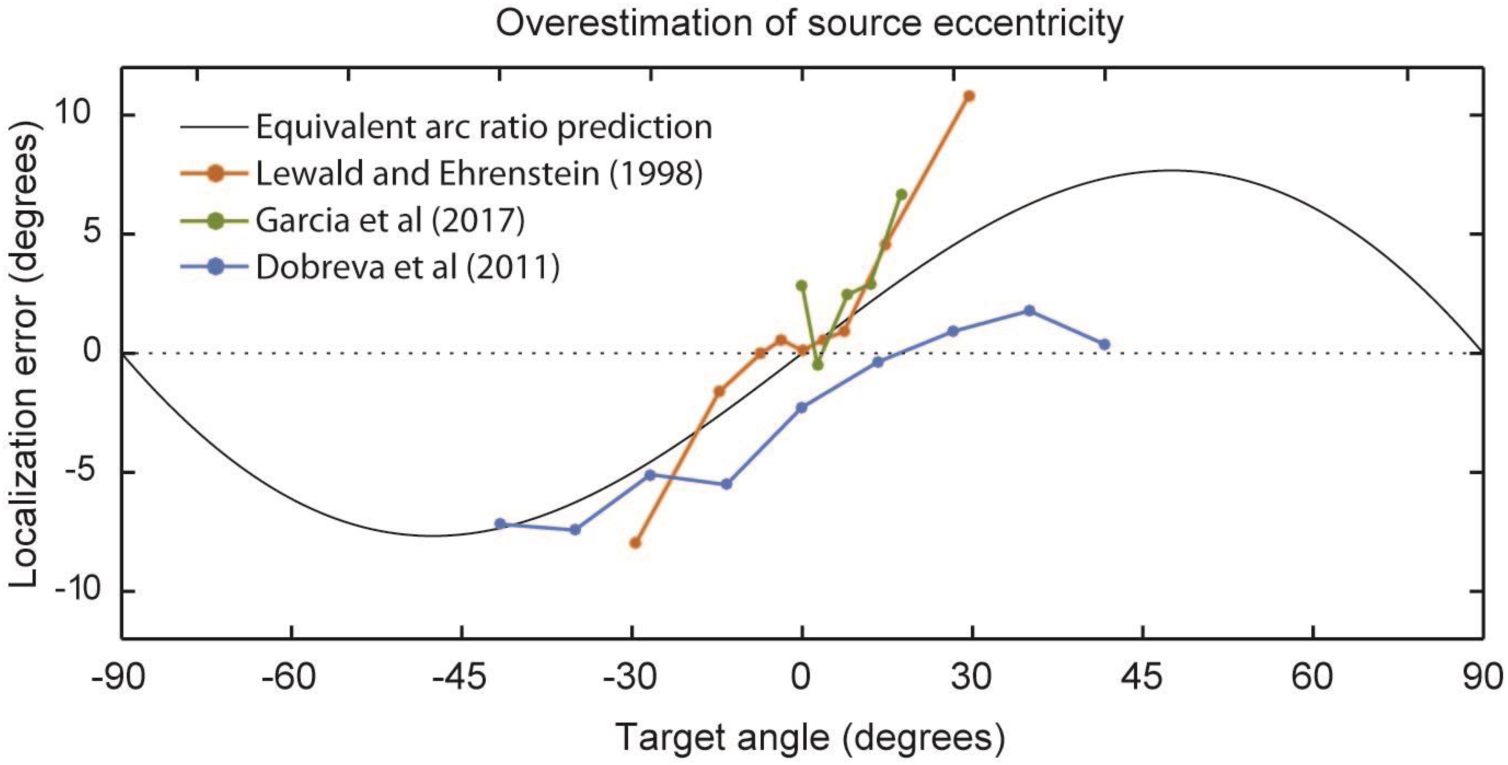
Predictions of overestimation of source eccentricity as a function of target angle. Data demonstrating that listeners overestimate target angles are displayed from three separate previous studies (colored dot symbols) alongside predictions of angle overestimation from Equation 1 (black line).

### 3.3 Distortion in acoustic motion

When Equation 1 is used to examine *motion* (by examining the differences in the distortion between *Θ*_a_ and *Θ*_p_ at different head and source angles) it becomes clear that the overestimation of static signal angle must move with the head. The consequence of this is that signals appear to move in different ways depending on their subtended angle with respect to the head during a turn. Figure 4 displays the way in which the apparent location of two sound sources (here a bird and a television) should shift as the listener turns to the right. A signal at 0° should shift to the left and a signal at 90° should shift to the right. Supplemental Figure S1 is an animation of this phenomenon depicting the perceived locations of 32 static signals arranged around the head as it turns. The angle of the listener’s nose is depicted as a line along the radius of the circle. The expansion/contraction in Figure 4 and in the animation is exaggerated by a factor of 2 for clarity.

**Figure 4.**
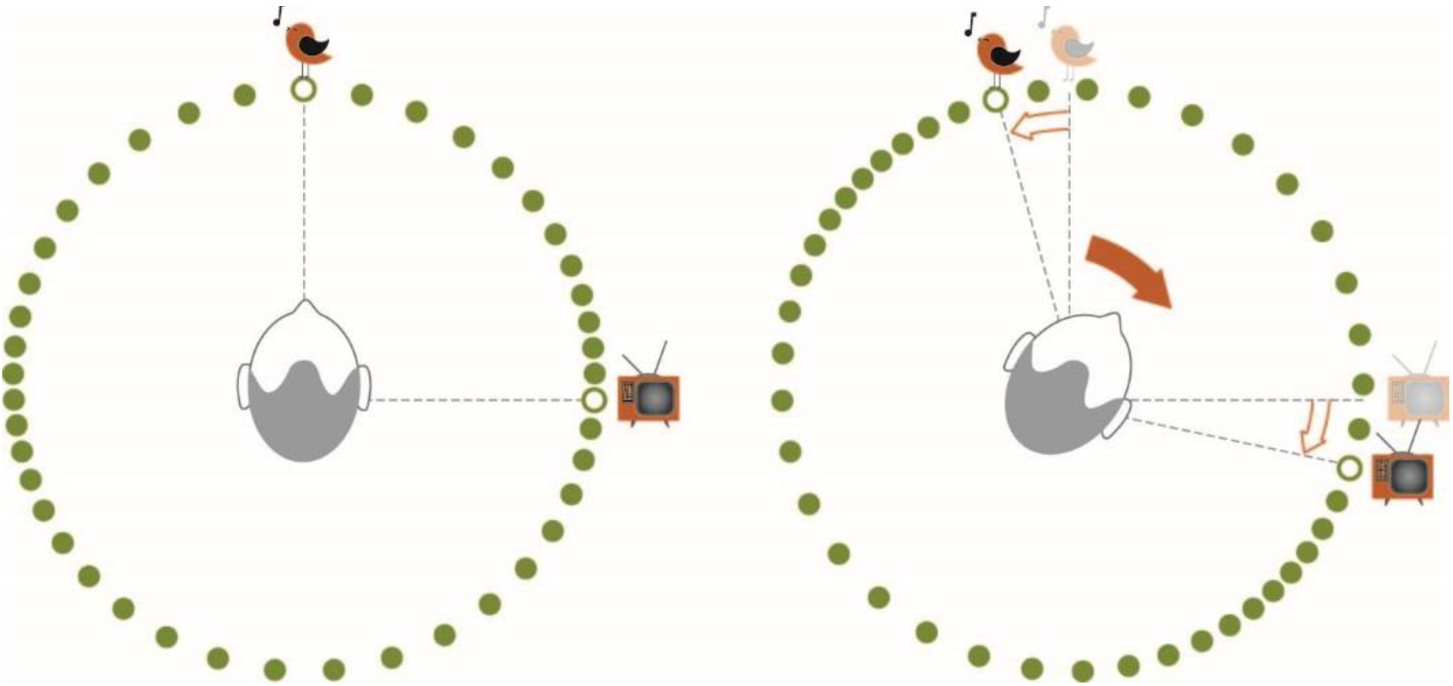
Illustration of the spatial distortion introduced by the equivalent arc ratio. The dots represent the perceived locations of 32 static signals arranged around the head as it turns to the right. The apparent location of a signal at the front moves leftward, whereas a signal at the right should appear move further to the right. The expansion represented here is exaggerated by a factor of 2 for the purpose of more clearly illustrating the phenomenon.

There are established phenomena that suggest there are perceptual distortions of auditory space that depend on some interaction between stimulus angle and head angle. Genzel and colleagues [7] demonstrated that, after an active head movement, a second sound source had to be shifted in azimuth to be perceived as being at the same azimuth as a sound before the movement. Within the framework of the equivalent arc ratio, this may be explainable as a distortion in the perceived location of a static midline signal. Using Equation 1 (with a 90°/0° PSE ratio of the mean 1.82), a 35.3° active rightward movement (the average reported in the study) should result in a 0° signal appearing to be at -6.5°, which not only sign-correct, it is also reasonably close to the value of -5.5° from Genzel et al (2016). Other systematic errors in movement compensation have also been documented. For example, Freeman et al [6] demonstrated that signals at the front of the listener must be moved with the head with a gain of +0.17 to be judged as being static. Here gain refers to amount of motion with respect to the head, so if a listener turns 10° to the right, signals that move by +1.7° to the right would be most consistently judged to be static. The corresponding value predicted by Equation 1 is +2.2°, which is at least sign-correct if not a perfect match.

Physiological data on the relationship between self motion and spatial receptive fields is virtually non-existent, making comparisons with animal work problematic. Eye position has been shown to clearly influence the apparent spatial location of auditory signals [19, 20], to modulate responses in the inferior colliculus [21] and auditory cortex [22], and to actively shift spatial receptive fields in superior colliculus [23], but little work has been done on head movements. Very recently, however, experiments in ferret primary auditory cortex have revealed a subpopulation of neurons whose spatial receptive fields appear to be specified in world-centric coordinates, rotated in opposition to the animal’s movement [24]. This finding represents a neural correlate of our percept of a stable acoustic world. It is not currently possible, however to determine whether the shifts in receptive field boundaries as a function of eccentricity match that predicted by the equivalent arc ratio because the width and contralateral offset of cortical receptive fields make it difficult to assign individual neurons to exact azimuths in space.

Returning to psychophysics work, results from the Freeman et al (2016) study are roughly line with what the equivalent arc ratio model would predict, with one notable exception. According to the model, the gain at which signals must be moved to be judged as static should change as a function of azimuth, reducing to 0 when 45° is reached, and even changing to a *counter*-rotation for larger eccentricities. The Freeman et al study did not find this, although they did find a decrease in gain and a substantial increase in variance as a function of stimulus angle. It should be noted that these authors’ own Bayesian explanation for the non-unity gain *also* predicts a change in gain as a function of azimuth. But the subjects in that study were blindfolded, so the discrepancy may point to an as-yet unresolved role of eye position in this and related phenomena. The previously mentioned dependence of neural and behavioral responses on eye position certainly attest to this possibility. A related phenomenon was previously described by Lewald and Ehrenstein [5]; the displacement of the subjective auditory midline (as measured using ILD) towards the trunk as a listener turns to more eccentric head angles. This displacement was argued as being the result of an ‘incomplete coordinate transformation,’ a failure of listeners to fully compensate for their own movement. Taken together with the results from Freeman et al [6] this suggests that head-to-trunk angle may represent a second unresolved factor that results in a shift in target location into a different region of expansion / contraction of acoustic space.

Studies examining representational momentum have argued that the faster a signal is rotating around the head, the further the perceived endpoint will be displaced in space [e.g., 25, 26]. This is argued to be a consequence of a mental extrapolation of the signal’s trajectory [27]. According to the equivalent arc model, signals moving towards the midline would appear to accelerate, suggesting their endpoints could seem more displaced than those of signals receding from the midline. An advantage in direction discrimination has been demonstrated for signals approaching the median plane [28], congruent with the equivalent arc model, but in the case of the first representational motion study [25] all the motion trajectories used were across the midline. The analysis in the second study [26] – while it did examine left versus right movements – collapsed the data across different center azimuths, an averaging method that would prevent us from observing any asymmetry predicted by the equivalent arc ratio model. Examination of the latter data set could either lend support to or require a re-evaluation of the equivalent arc framework.

### 3.5 The relationship between the equivalent arc ratio and the MAMA

The equivalent arc ratio expansion observed appears to be related to – but not entirely dependent on – the change in MAMA as a function of angle (the MAMA being roughly 1° in front of the listener and increasing to about 4° at the side [4]). If the equivalent arc ratio were a simply the result of the change in MAMA as a function of angle, then we might expect slightly larger equivalent arc ratios between 0° and 90° than we observed. However, the two measurements may be linked with each other on some level, as acoustic movement, whether a consequence of source or self motion, may rely on similar underlying processing mechanisms [c.f. 29]. We did not test the MAMA at 0°, 45°, and 90° in our listeners, so we cannot at this point describe the correlation between the two measures.

### 3.6 Creating hyper-stable virtual acoustics

Because listeners may perceive signals to move at different velocities at different points in the arc around the head, the equivalent arc ratio could be utilized alongside individualized head related transfer functions and motion tracking to produce head-stabilized acoustic environments that appear to be more stable than the real world. As seen from the scatter in Figure 2, the PSE ratio can vary greatly from listener to listener. As such this must be measured or approximated through other means to match a given listener’s spatial distortion. Given the close relationship between the equivalent arc predictions and previously described overestimations of target angle, it may be sufficient simply to have a listener point to a few sound sources with a laser. Regardless of how this is measured, an inverse of Equation 1 that is solved for *Θ*_a_ would be necessary. This is included here as Equation 2.

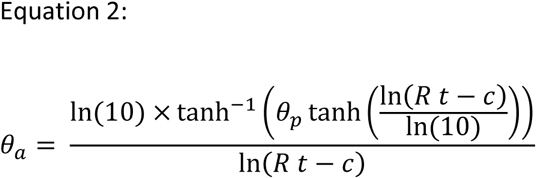

where the constants and definitions in the formula are the same as in Equation 1.

This formula allows one to determine the angles at a signal must be presented to be perceived at a particular azimuth with respect to the head.

### 3.7 Caveats

The range of movement excursions in this study was not sufficient to compare references at 0° and test signals at 90° for all listeners. We did not anticipate the magnitude of the spatial expansion that we observed and so were not able to fully bracket the motion values and measure PSE ratios for all movement pairs. We were able to measure PSE ratios for the inverse of these particular reference/test pairs, but direct comparison between these makes the tacit assumption that the amount of spatial expansion/contraction is a simply a multiple of the reference motion.

It should be also noted that Equation 1, while it may be reasonably applicable to perceptual distortion of signal location in the listener’s front hemifield they may not accurately reflect any expansion or contraction of auditory space in the *rear* hemifield (and indeed Equation 1 is not constructed to compute the perceived location of angles beyond ±90°). We have no data that speaks to this, so the expansion and contraction in the rear hemifield is depicted in Figure 4 and Supplemental Figure S1 as a mirror reflection of the front, despite there being no reason to believe this is necessarily the case. It remains for future studies to map out spatial distortions for 360° around the head.

More generally speaking, since expansion estimates were only measured at three angles, it is unclear whether a hyperbolic tangent expansion or some other function may be the most appropriate mathematical descriptor of the change in equivalent arc ratio over all azimuths. Future work will be required to determine what function best captures the observed phenomena but – provided the function is readily invertible – such a technique could potentially increase the experience of immersion for virtual reality systems.

### 3.8 Conclusions

Sound sources at the side of a listener must move at least twice as much as ones in front to be judged as moving the same amount. This expansion of space in the front and compression at the side that moves with the listener we term the equivalent arc ratio, and likely has real consequences for spatial perception in dynamic listening situations. The prediction that the apparent location of static sound sources may also be distorted suggests that this phenomenon is not limited to moving signals. A mathematical model that mimics the equivalent arc ratio can be used to successfully predict several previously unexplained phenomena in spatial auditory perception. We further suggest that the inverse of this function could be utilized alongside individualized head related transfer functions and motion tracking to produce head-stabilized virtual acoustic environments that appear to be more stable than the real world.

## 4 Materials and Methods

### 4.1 Participants

We recruited 30 normal-hearing listeners, with normal hearing being defined as a four-frequency average pure tone hearing threshold of less than 20 dB HL. Five listeners were excluded from the analysis because they did not complete the full set of trials. We collected complete data sets for the remaining 25 listeners, the result of two separate visits to the lab, with sessions of 60 minutes each. The average age of the listeners was 27 (±7.3 STD) years, ranging from 22 to 58 years old. We received written and verbal informed consent from all subjects and the experiment was conducted in accordance with procedures approved by the West of Scotland Research Ethics Service.

### 4.2 Stimuli and Presentation

The experiment was conducted in a 4.8 × 3.9 × 2.75 m double walled, sound-attenuated chamber that had 10 cm acoustic wedge foam lining the walls and ceiling, but not the floor, which was carpeted. The listeners were seated in this chamber in the center of a 3.5 m diameter circular ring of 24 Tannoy VX-6 loudspeakers (Tannoy, Coatbridge, UK) placed at intervals of 15°. Because a forward (towards the 0° loudspeaker) offset in listener position could yield an apparent expansion in space, the listener’s head was aligned with a spot on the ceiling and the floor. This method, while subject to a few centimeters of error, prevented a misplacement that could explain the results observed (which would require the listener to be at least an order of magnitude closer to the front loudspeaker). The room was dimly lit, but the loudspeakers were visible, and listeners were asked to keep their head still and their eyes open and fixated on the loudspeaker ahead of them at 0°. Signal sources were moved around the ring using vector-based amplitude panning (performed on a sample-by-sample basis in Matlab 2015b (The Mathworks, Natick, MA, USA) using the open source dynamic link library “playrec” (<u>www.playrec.co.uk)</u>). The signals were played out using a MOTU 24 I/O (Mark of the Unicorn, Cambridge, MA, USA) over ART SLA-4 amplifiers (Applied Research & Technology ProAudio, Niagara Falls, NY, USA). The stimuli were unfrozen pink noise signals that were amplitude modulated with a 10 Hz reverse sawtooth waveform at a depth of 50%. These signals provided sufficient high frequency energy to provide robust interaural level differences as well as frequent sharp onset transients to ensure that the signals were easily localizable. All signals were presented at a comfortable listener-determined listening level (this ended up being between 70 and 75 dB SPL).

### 4.3 Experimental Paradigm

We measured the point of subjective equality (PSE) for amount of acoustic motion between “test” and “reference” signals. The reference signal always moved 20° in a random direction and the test signal moved either less, the same, or more (5, 10, 15, 20, 25, 30, or 35°), also in a random direction (see Figure 5). The order of the test and reference signals was also randomized. In a two-alternative forced choice paradigm, listeners were asked to report on a touchscreen whether the first or second signal ‘moved more.’ If the listeners requested clarification on these instructions, they were told that their task was to report whether the first or second noise moved over a larger distance in space, regardless of its duration or apparent speed. Both the reference and the test signals could be centred at either 0°, 45°, or 90° (See Figure 5) plus or minus a random value drawn from a uniform distribution between -7.5 and 7.5°. The duration of the test and reference signals were individually randomized on every trial to a value between 0.5 and 2 seconds. In this way, we mitigated velocity and duration as potential cues, leaving total angular excursion as the variable that listeners were asked to judge. Listeners were asked to complete a total of 10 blocks of 126 trials, each of which contained 6 repeats of the 21 conditions.

**Figure 5.**
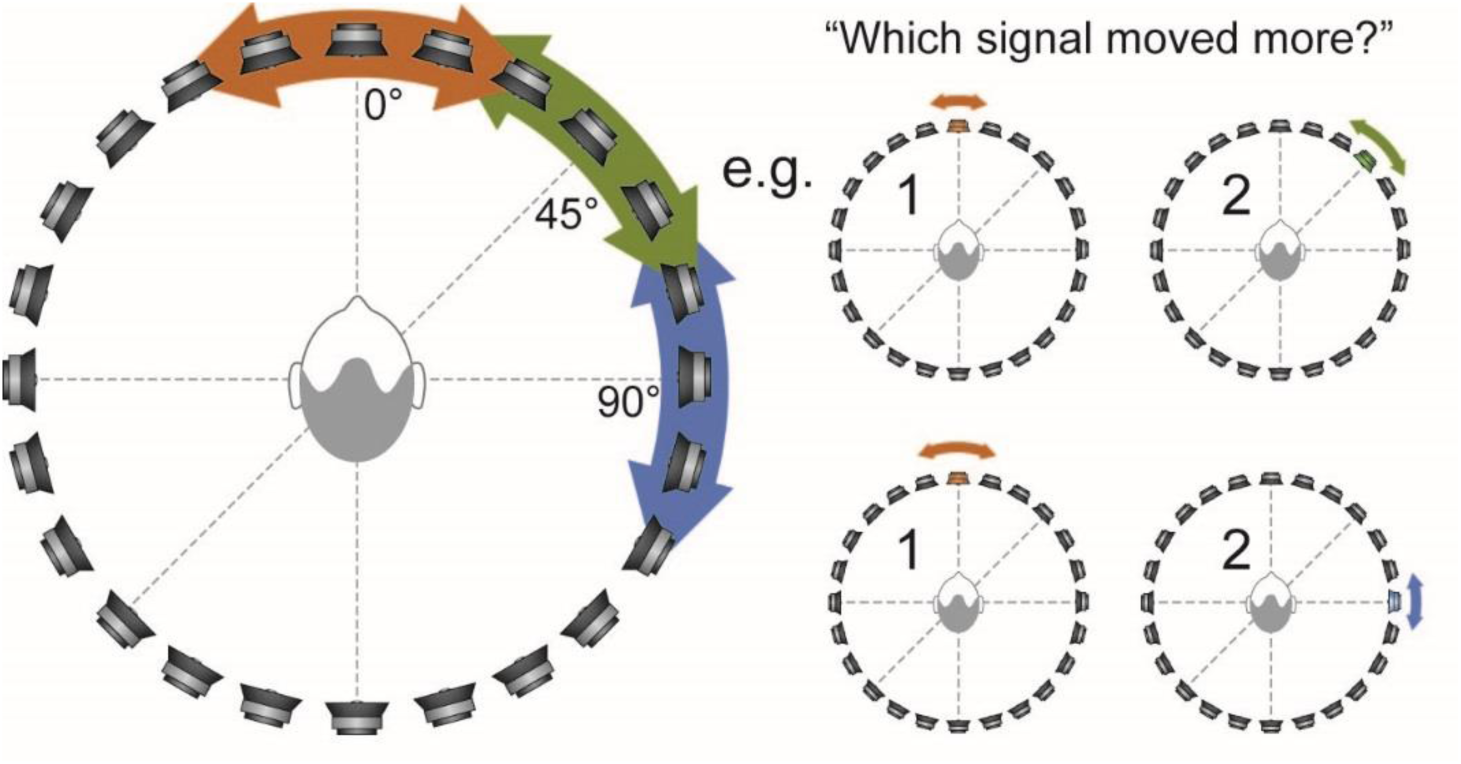
Experimental Paradigm: Listeners were presented with moving reference (20°) and test signals (variable °) at three possible center angles (0°, 45°, and 90°), randomized in order, and asked to report which of the two signals moved more.

The resulting psychometric functions for each listener were individually fitted with a logistic function using Matlab’s fminsearch function. The resulting parameters were fed into an inverse logistic equation to compute the test excursion value at which the function crossed the PSE (the reader may roughly infer these values in Figure 1 as the point where the mean fit (dotted line) crosses 0.5 on the y-axis). For logistic fits that did not cross the PSE before 40° (the next larger measurement point step) we fixed the value at 40°. This likely underestimates the true PSE for many listeners, but avoids excessive extrapolation. This value was divided by the reference excursion of 20° to yield a ratio expressing the amount of expansion or contraction of auditory space.

### 4.4 Statistics

All statistics were performed with the Statistics Toolbox in Matlab 2016a. The analysis consisted of a three-way repeated measures ANOVA with the dependent variable being the proportion of ‘moved more’ responses, and the independent variables being reference angle, test angle, and test excursion. Alpha was set to 0.05.

## Acknowledgements

Thanks to Kay Wright-Whyte and Jack Holman for help collecting data and Graham Naylor and Alan Archer-Boyd for reading drafts of this manuscript.

---

**Supplemental Figure S1.**
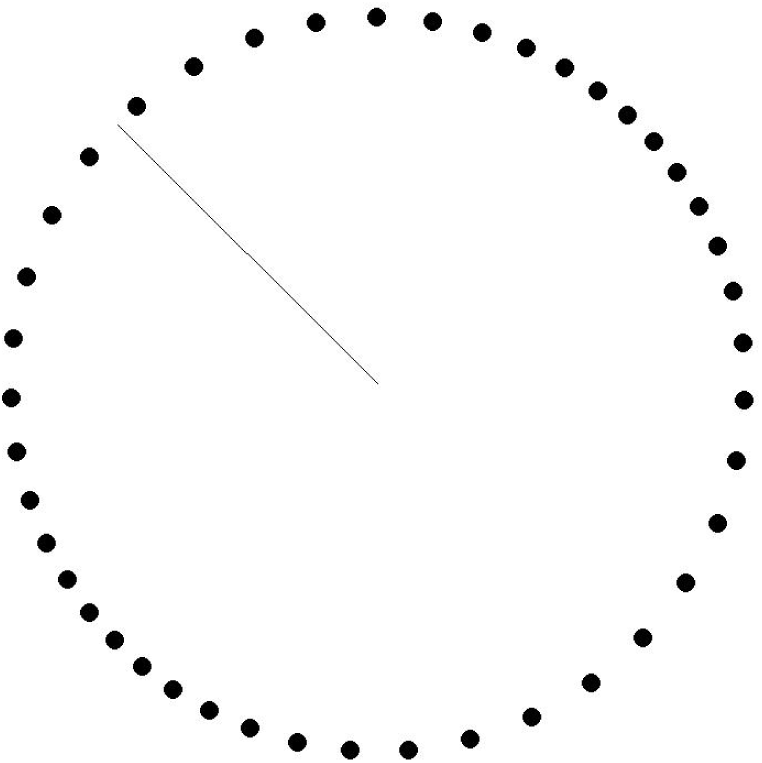
Animation of the spatial distortion introduced by the equivalent arc ratio. The dots represent the perceived locations of 32 static signals arranged around the head as it turns. The angle of the head is represented by the line along the radius of the circle. The expansion represented here is exaggerated by a factor of 2 for the purpose of more clearly illustrating the phenomenon.

